# T-bet expressing Tr1 cells driven by dietary signals dominate the small intestinal immune landscape

**DOI:** 10.1101/2025.06.30.662190

**Authors:** Eduard Ansaldo, Daniel Yong, Nathan Carrillo, Taryn McFadden, Mahnoor Abid, Dan Corral, Claudia Rivera, Taylor Farley, Nicolas Bouladoux, Inta Gribonika, Yasmine Belkaid

## Abstract

Intestinal immunity defends against enteric pathogens, mediates symbiotic relationships with the resident microbiota, and provides tolerance to food antigens, safeguarding critical nutrient absorption and barrier functions of this mucosal tissue. Despite the abundance of tissue resident activated T cells, their contributions to these various roles remains poorly understood. Here, we identify a dominant population of IL-10 producing, T-bet expressing CD4+ Tr1 T cells, residing in the small intestinal lamina propria at homeostasis. Remarkably, these intestinal Tr1 cells emerge at the time of weaning and accumulate independently of the microbiota displaying similar abundance, function and TCR repertoire under germ-free conditions. Instead, the small intestinal T-bet+ Tr1 program is driven and shaped by dietary antigens, and accumulates in a cDC1-IL-27 dependent manner. Upon activation, these cells robustly express IL-10 and multiple inhibitory receptors, establishing a distinct suppressive profile. Altogether, this work uncovers a previously unappreciated dominant player in homeostatic small intestinal immunity with the potential to play critical suppressive roles in this tissue, raising important implications for the understanding of immune regulation in the intestine.

**Significance Statement:** Establishing immunological tolerance to self and environmental antigens is critical to preserve tissue homeostasis and function. In the intestine, both dietary and microbiota derived antigens are routinely encountered by the immune system, which deploys a variety of mechanisms to maintain tolerance to these innocuous antigens. Understanding how immunological tolerance is established is critical, a when this process goes awry it can lead to severe inflammatory and autoimmune diseases such as food allergy and inflammatory bowel disease. However, how tolerance is established in the intestine is still poorly understood. In this study we describe a novel dominant T cell population in the small intestine shaped by dietary components with the potential to play important roles in immune tolerance at this site. back # Introduction

Barrier surfaces such as the gut and skin represent the first line of defense against the environment. These organs must strike a delicate balance between providing protection against environmental and infectious agents, maintaining tissue function, and establishing a homeostatic symbiotic relationship with resident microbes collectively known as the microbiota (1). The immune system plays a critical role in establishing these dynamic and carefully regulated relationships, as evidenced by the large number of immune cells present at these sites. Of particular note, activated T cells are very abundant at barrier tissues, where they orchestrate immune effector functions geared towards these varied tasks (1, 2). In the small intestine, the intraepithelial compartment harbors innate like natural CD8aa⁺ IELs, many of which are self reactive; as well as CD4⁺CD8aa⁺ and CD8ab⁺ IELs responding to dietary and microbial antigens (3). The underlying lamina propria (SILP) harbors predominantly CD4⁺ T cells, which participate in responses to commensal-derived and dietary antigens (2, 4). Despite the abundance of small intestinal CD4 T cells, only a handful of cognate immune interactions focusing on Type 17 and T regulatory helper subsets have been described. Thus, whether immune responses in this tissue are truly limited to a small number of antigenic triggers and effector functions remains to be fully elucidated.

The small number of gut homeostatic CD4 T cell responses described thus far have been shown to primarily respond to specific commensal bacteria or dietary antigens (1, 2, 5–8): Among other examples, SFB induces cognate Th17 cells in the small intestine (9, 10), a consortium human commensal bacteria induces CD8b⁺ cells in the colon (11), and *Akkermansia muciniphila* indices T_FH_ and other effector cells in the Peyer’s patches and lamina propria, respectively (12). Furthermore most regulatory T cells in the colon are induced in response to commensal or pathobiont species at homeostasis, providing critical regulatory functions (13, 14). Cognate immune responses to SFB help contain this commensal species in the intestine (15), but also have systemic impacts on the susceptibility to autoimmune disease (16, 17). Interestingly, despite presenting a classical Th17 effector profile, a subset of SFB-induced Th17 cells possess IL-10 secretion capabilities and suppress cognate immune responses without the expression of Foxp3 (18), suggesting immunoregulatory functions reminiscent of Tr1 cells. Whether these competing capabilities are unique to SFB-specific immune responses or a general hallmark of small intestinal immunity remains unknown.

The description of SFB-specific Tr1-like cells in the small intestine was surprising, as this CD4⁺ T cell subset, characterized by abundant IL-10 secretion in the absence of *Foxp3* expression, has only been described in the context of chronic antigen stimulation, such as chronic infection or cancer (19). The Tr1 cell program is controlled by a variety of transcription factors and upstream signaling pathways, including IL-27 signaling, MAF and AHR (20). AHR-ligands are abundant in the intestine, and MAF is a hallmark of other regulatory commensal-specific responses (21, 14). Furthermore, IL-27, which can induce both proinflammatory and immunoregulatory functions, is abundant in the small intestine (22, 23). This raises the possibility that the Tr1 program is a more general feature of small intestinal immunity, not uniquely restricted to SFB-specific responses.

In this study we explore the breadth of CD4⁺ T cell responses in the small intestine, and uncover a previously uncharacterized CD4⁺T-bet⁺ T cell immune response that is dominant in this tissue. Unexpectedly, these SILP CD4⁺T-bet⁺ T cells are independent of the microbiota, maintaining a similar functional profile and shared antigen specificities in germ-free conditions. Instead, we reveal that dietary components drive the accumulation, function, and clonal selection of this T cell population. Finally, we show that, contrary to classical Th1 cells, SILP CD4⁺T-bet⁺ T cells adopt a Tr1 immunoregulatory functional program during activation, suggesting that this is a general feature of CD4⁺ T cell immunity in the small intestine wired towards immune regulation and tissue homeostasis.

## Results

### The small intestinal immune landscape is dominated by CD4⁺T-bet⁺ cells

We profiled the small intestinal (SI) T cell compartment in mice by flow cytometry and immunofluorescence using a combination of transcription factor staining and reporter mice (Figures 1A-E). Surprisingly, in contrast to established views in the literature (1, 2), T-bet expressing lymphocytes, in particular CD4⁺T-bet⁺Foxp3⁻ T cells (herein called CD4⁺T-bet⁺ T cells), dominated the T cell compartment in the small intestine lamina propria, outnumbering the well characterized SI Th17 and Treg cells, and comprising almost 50% of all T lymphocytes in this tissue (Figures 1A-C). CD4⁺T-bet⁺ T cells comprised ∼50 percent of CD4⁺ T cells in this compartment, compared to 25 and 10 percent of Th17 and Treg cells, respectively (Figure 1A-B). Confocal imaging of T-bet expression confirmed the presence of T-bet expressing T cells across the small intestine (Figure 1D-E), with CD4⁺T-bet⁺ T cells detected in the villi and crypts in the tissue underlying the epithelium, known as lamina propria (Figure 1D). CD4⁺T-bet⁺ T cells were most abundant in the small intestine lamina propria, while also detected in the intraepithelial compartment of the small intestine but with very low numbers detected in the large intestine (Figure 1F). Interestingly, CD4⁺T-bet⁺ T cells were more abundant proximally in the duodenum, compared to the distal small intestine (Figure 1G).

**Figure 1:**
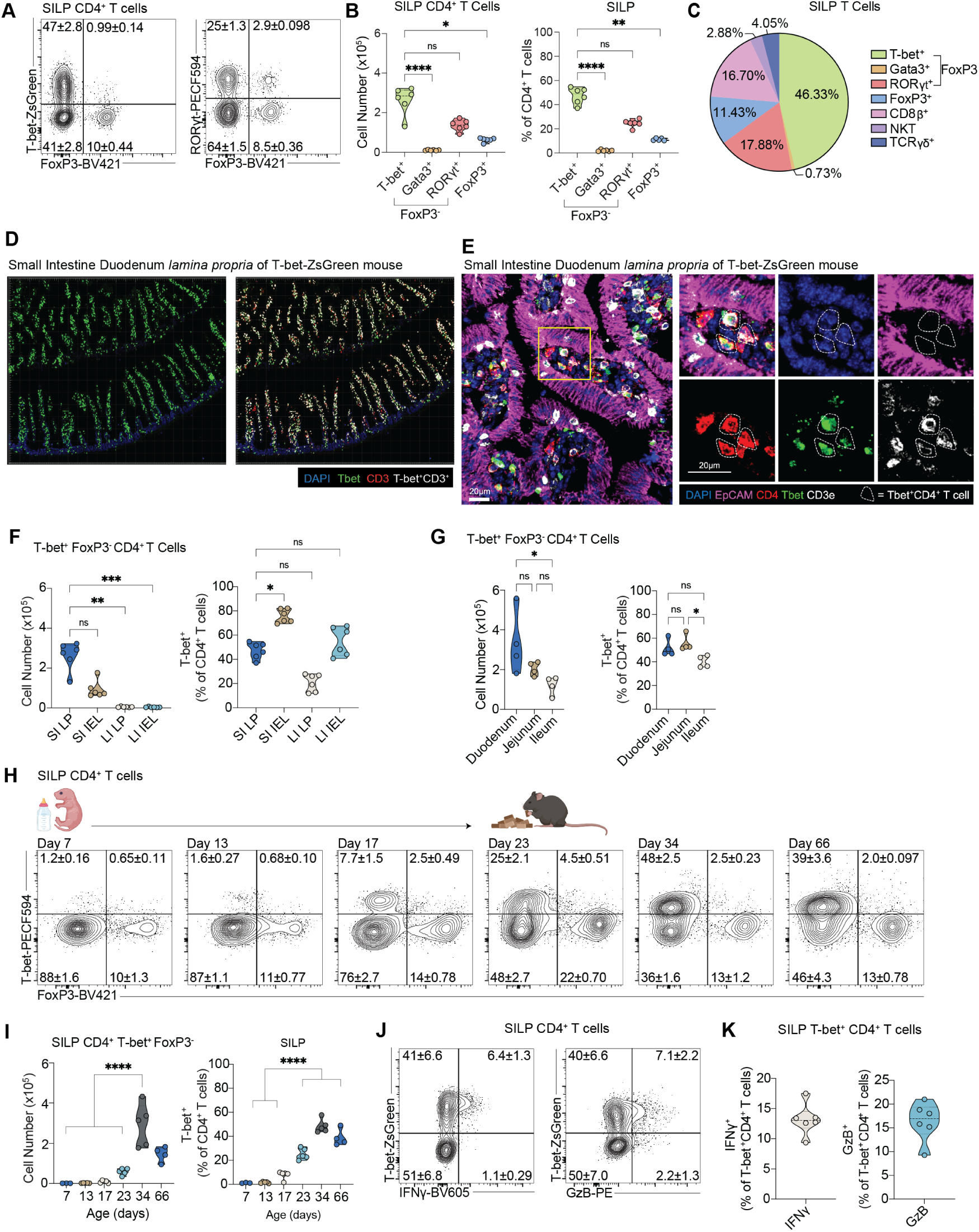
The small intestinal immune landscape is dominated by CD4⁺T-bet⁺ T cells. (A - B) Representative flow cytometry plots showing SILP T-bet+ and RORgt+ frequency on gated CD45+ CD90.2+ TCRb+ CD4+ T cells (A) and violin plots showing T-helper subsets including T-bet+, Gata-3+, RORgt+, and FoxP3+ cell counts and frequencies (B). (C) Relative proportion of T cell subsets isolated from SILP as measured by flow cytometric analysis. (D - E) Confocal images of the small intestine lamina propria of SPF T-bet-ZsGreen reporter mice at steady state. (F) Violin plots showing SILP T-bet+ cell count and frequencies in the lamina propia and intraepithelial compartment in the small and large intestine. (G) Violin plots showing SILP T-bet+ cell count and frequencies across different small intestinal sections (duodenum, jejunum, and ileum) (H - I) Representative flow plots showing kinetic of SILP Tbet+ frequency from neonatal to adulthood on gated CD45+ CD90.2+ TCRB+ CD4+ T cells (H) and violin plots showing SILP T-bet+ cell counts and frequencies (I). (F, G, I) Gated on CD45+ CD90.2+ TCRb+ T-bet+ FoxP3- CD4+ T cells. Data are representative of at least three independent experiments. Ns, not significant, *P < 0.0332, **P < 0.0021, ***P < 0.0002, ****P < 0.0001. Kruskal-Wallis test. Error bars indicate SEM.

The switch from milk to solid food represents a critical milestone in the development and maturation of the intestinal immune system (24, 1, 25, 26). This transition is characterized by a profound remodeling of the intestinal microbiota driven by the shift in dietary composition and complexity (27). Concomitantly, the intestinal immune compartment undergoes rapid development and maturation during this period, with an acute influx and expansion of lymphocyte populations (28). Similarly, we observed rapid accumulation of CD4⁺T-bet⁺ T cells between the third and fifth week of life (Figure 1H-I), which corresponds to the weaning period in mice. These data suggest that CD4⁺T-bet⁺ T cells also accumulate in response to dietary and/or microbiota derived signals. Interestingly, only a fraction of SILP CD4⁺T-bet⁺ T cells presented with classical Type-1 effector functions such as IFN𝛾 and TNF cytokine production or GranzymeB expression (Figure 1J-K, S1A-B), suggesting that other capabilities beyond classical Th1 effector functions may be present in this population. Finally, we tested whether SILP CD4⁺T-bet⁺ T cells depend on classical MHC-II antigen presentation. *I-Ab^-/-^* mice presented with an almost complete lack of CD4⁺T-bet⁺ T cells in the small intestine lamina propria (Figure S1C-E), consistent with restriction on MHC-II and a conventional CD4 T cell origin.

Together, these data uncover CD4⁺T-bet⁺ T cells as a dominant component of homeostatic T cell immunity in the small intestine, and support the idea that signals from dietary components or the microbiota may drive the accumulation of this population.

### Small Intestinal CD4⁺T-bet⁺ T cells are independent of the microbiota

Next, we interrogated whether SILP CD4⁺T-bet⁺ T cells are driven by microbiota derived antigens. To that end we rederived T-bet-ZsGreen mice into germ-free conditions and examined the accumulation of this population in the absence of microbes. Interestingly, germ-free mice presented similar levels of SILP CD4⁺T-bet⁺ T cells at homeostasis as compared to SPF mice with similar functional capacity (Figure 2A.B and S2A-F), in contrast to Th17 cells, which are known to depend on the microbiota (9, 10).

**Figure 2:**
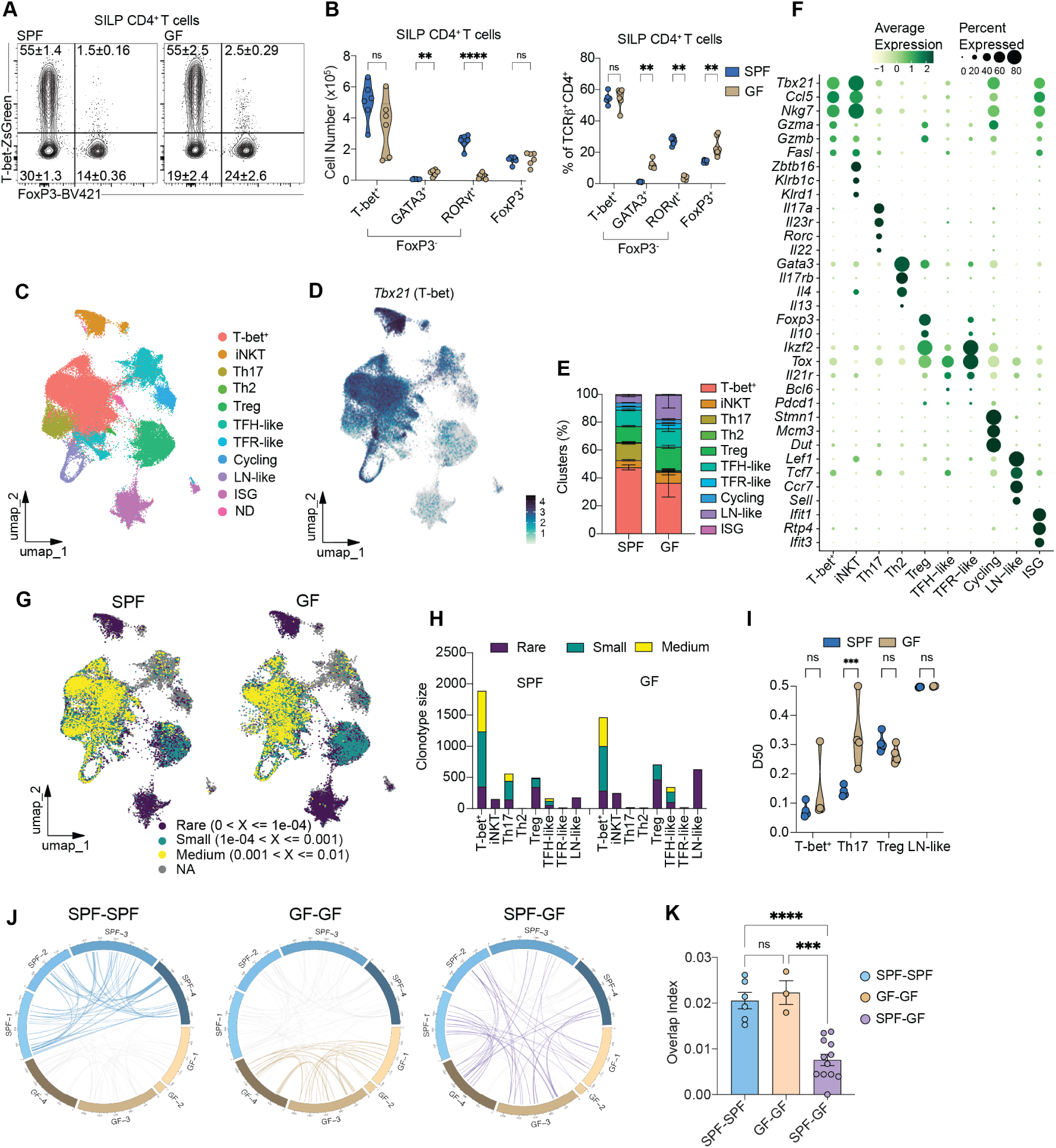
Small intestinal CD4⁺T-bet⁺ T cells are independent of the microbiota. (A - B) Representative flow plots showing SILP CD4+ frequency from SILP of SPF and GF mice (A) and violin plots showing T-helper cell counts and frequencies between SPF and GF mice (B). (C-J) Single Cell RNA-seq and VDJ sequencing of SILP CD4 T cells from SPF and germ free mice (4 mice per group). (C) Uniform Manifold Approximation and Projection (UMAP) plot of SILP CD4 T cells. Clusters are annotated into 11 cell types. (D) Single cell expression of *Tbx21* (T-bet) projected on the UMAP plot of SILP CD4 T cells. Cells where *Tbx21* expression is not detected are shown in gray. (E) Bar graph representing the cell type relative frequency per condition of the clusters represented in (C). (F) Dot plot showing the normalized expression of the indicated genes in each cluster. Dot size represents the faction of the cells where the gene is detected, and the color scale represents row z-scored average normalized expression. (G) T cell receptor clonotype size distribution per sample projected onto the UMAP plot of SILP CD4 T cells split by condition. Data are quantified on a bar graph in (H). Medium: clonotypes that represent between 0.1% and 1% of all TCR sequences. Small: clonotypes that represent between 0.01% and 0.1% of all TCR sequences. Rare: clonotypes that represent less than 0.01% of all TCR sequences. (I) D50 clonotype diversity measure of the indicated clusters per condition. (J) Circos plots highlighting clonotypes shared within SPF mice (left), within GF nice (center), or between SPF and GF mice (right). Each section represents one mouse, and each link represents a shared TCR clonotype, with the width of the link at each end representing the abundance of that TCR clonotype in that sample. (K) Quantification of the TCR repertoire overlap in (J). Data are representative of at least three independent experiments (A -B). Ns, not significant, *P < 0.0332, **P < 0.0021, ***P < 0.0002, ****P < 0.0001. Multiple unpaired t tests with Welch correction (A, I), One-way ANOVA (K). Error bars indicate SEM.

To further characterize the CD4 T cell compartment in the small intestine lamina propria, we performed single cell RNA sequencing (scRNAseq) of small intestine lamina propria CD4⁺ T cells at homeostasis in germ-free and SPF conditions. We performed unsupervised clustering and annotation of the resulting clusters into cell types (See methods). With this, we were able to identify all major T helper subsets based on cell type specific markers (Figure 2C-F). Similar to our flow cytometry approach, we observed a T-bet⁺ T cell cluster in similar proportion in both germ-free and SPF mice, while the Th17 cluster was absent in germ-free mice (Figure 2E, S2G-H). Interestingly, we observed a small distinct T-bet⁺ cluster corresponding to CD4⁺ invariant NK cells (iNKT), which we annotated based on expression of PLZF, NK cell receptors, and the iNKT chain (Figure 2C, F, S2G, I). Importantly, the dominant T-bet⁺ population lacked expression of these factors (Figure 2F, S2I), clearly establishing the majority of CD4⁺T-bet⁺ cells as a distinct population from iNKTs. Thus, we subsequently excluded iNKT cells from the T-bet⁺ gate in our flow cytometry analysis using the iNKT CD1d tetramer. Finally, we further characterized this T cell population by exploring the expression of several factors associated with transcriptional regulation, tissue residency and migration, T cell activation, effector function, and co-stimulation (Figure S2J-L). Of note, T-bet⁺ cells expressed high levels of the chemoattractant *Ccl5* and the co-stimulatory receptor *Cd40lg*, which it shared with iNKTs and Th17 cells, respectively (Figure S2L). Furthermore, the transcription factors *Maf* and, to a lesser degree, *Ahr* were widely expressed across SILP CD4 T cells, excluding the lymph node-like cluster (Figure S2J-L). Finally, T-bet⁺ cells expressed many factors associated with tissue residency and migration in the small intestine (*Ccr9*, integrin), T cell activation (*Cd69*,*Cd44*,*Nr41*), cytotoxicity (*Gzma*, *Gzmb*, *Nkg7*), and co-stimulation (*Ctla4*, *Cd28*, *Cd40lg*), features which it shared with iNKTs, Th17, and/or Tregs in this tissue (Figure S2L).

In parallel, we characterized the T cell receptor repertoire of small intestinal lamina propria CD4⁺ T cells at the single cell level. This revealed the presence of a polyclonal repertoire with evidence of clonal expansion in CD4⁺T-bet⁺ T cells in both SPF and germ-free conditions, while Th17 cells were clonally expanded only in SPF but not germ-free mice (Figure 2G-I). These data reveal that the TCR repertoire of CD4⁺T-bet⁺ T cells is not controlled by the microbiota, in contrast to previously described homeostatic Th17 T cell populations. Finally, we profiled the TCR repertoire of SILP CD4⁺T-bet⁺ T cells for the presence of shared (public) clonotypes. Interestingly, we found a significant degree of clonal sharing between mice within both SPF and germ-free conditions (Figure 2J-K). Importantly, we also detected clonal sharing across SPF and germ-free mice, albeit at lower level than within conditions, suggesting shared antigen specificities of SILP CD4⁺T-bet⁺ T cells independent of microbiota status (Figure 2J-K).

These findings strongly support the argument that SILP CD4⁺T-bet⁺ T cells do not depend on or respond to microbiota derived signals or antigens, in contrast to previously described T helper subsets present at homeostasis in the intestine. This phenotype mirrors T regulatory cells in the small intestine, another T cell population that is largely independent of the microbiota at this site, instead responding to dietary antigens and providing oral tolerance (13, 7, 5). The independence of the microbiota raises the possibility that SILP CD4⁺T-bet⁺ T cells may instead be responding to dietary antigens.

### Dietary components drive the accumulation and clonal selection of small intestinal CD4⁺T-bet⁺ T cells

The induction of cognate regulatory T cell responses to dietary antigens is critical to maintain immune tolerance and preserve tissue homeostasis in the intestine (5–7). Recent work highlighted the control of regulatory cells and defined IEL populations by dietary antigens (7, 6, 8), however the extent to which diet controls other immune cell populations in the small intestine remains poorly understood. Indeed, given the low abundance of microbes in proximal regions of the small intestine, it is likely that other environmental signals such as diet may play a dominant role in immune system function. To test whether SILP CD4⁺T-bet⁺ T cells are controlled by dietary anti-gens, we designed two refined diets with low (casein diet) or no abundance of dietary polypeptides (aminoacid diet) (See Methods). Germ-free mice raised under casein or amino acid diet showed a stark decrease in CD4⁺T-bet⁺ T cells in the small intestine (Figure 3A-B), suggesting that dietary signals control the accumulation of this population and that a diverse dietary polypeptide repertoire may be driving cognate CD4⁺T-bet⁺ T cell responses.

**Figure 3:**
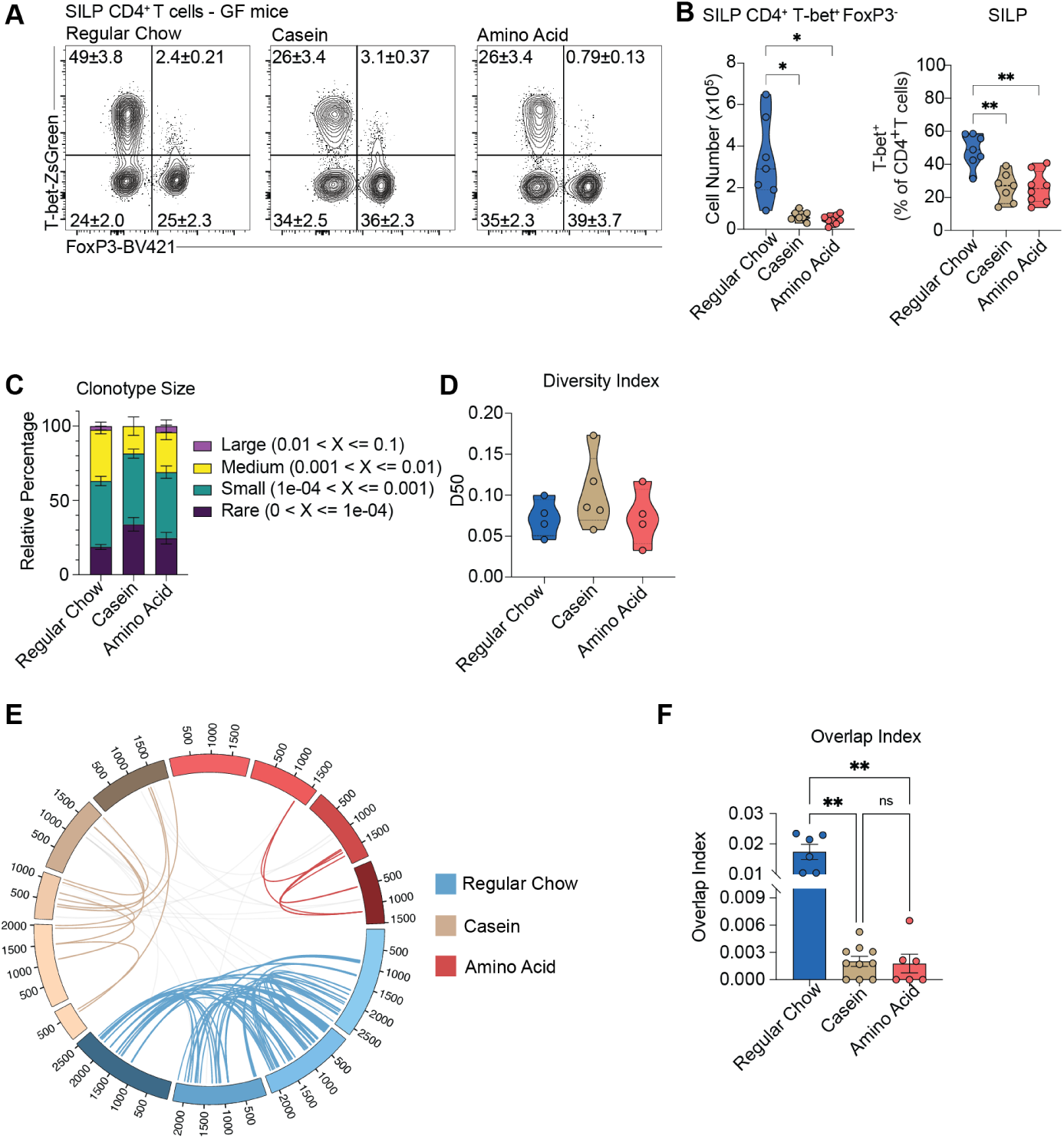
Dietary components drive the accumulation and clonal selection of small intestinal CD4⁺T-bet⁺ T cells. (A) Representative flow plots showing SILP Tbet+ frequency from SILP of GF mice fed either regular chow, casein, or amino acid diet since birth. (B) Violin plots showing SILP Tbet+ count and frequency on gated CD45+ CD90.2+ TCRb+ T-bet+ Foxp3-cells from (A) (C - F) Single Cell VDJ sequencing of SILP CD4⁺ T-bet⁺ FoxP3⁻ T cells from GF mice fed either regular chow, casein, or amino acid diet (4-5 mice per group). (C) T cell receptor clonotype size distribution per sample. Large: clonotypes that represent between 1% and 10% of all TCR sequences. Medium: clonotypes that represent between 0.1% and 1% of all TCR sequences. Small: clonotypes that represent between 0.01% and 0.1% of all TCR sequences. Rare: clonotypes that represent less than 0.01% of all TCR sequences. (D) D50 clonotype diversity measure of SILP CD4⁺ T-bet⁺ FoxP3⁻ T cells per condition. (E) Circos plots highlighting clonotypes of SILP CD4⁺ T-bet⁺ FoxP3⁻ T cells shared within GF regular chow-fed mice, casein diet-fed mice, or amino acid diet-fed mice. Each section represents one mouse, and each link represents a shared TCR clonotype, with the width of the link at each end representing the abundance of that TCR clonotype in that sample. (F) Quantification of the TCR repertoire overlap index in (E). Data are representative of at least three experiments (A - B). Ns, not significant, *P < 0.0332, **P < 0.0021, ***P < 0.0002, ****P < 0.0001. Brown-Forsythe and Welch ANOVA tests. Error bars indicate SEM.

To further understand the impact of diet on the TCR repertoire of CD4⁺T-bet⁺ T cells, we performed single cell RNA and TCR sequencing on equal numbers of sorted SILP CD4⁺T-bet⁺ T cells from germ-free mice on regular chow, casein, or amino acid diet. While the degree of clonal expansion was not impacted on the remaining SILP CD4⁺T-bet⁺ T cells (Figure 3C-D), the large degree of clonal sharing between mice raised on regular chow was almost completely absent when mice were raised on a casein or amino acid diet (Figure 3E-F). These data reveal that diet controls the accumulation and the clonal selection of CD4⁺T-bet⁺ T cells in the small intestine.

### cDC1s and IL-27, but not IL-12, are required for the accumulation of SILP CD4⁺T-bet⁺ T cells

T-bet expressing CD4⁺ T cells are typically associated with Th1 responses, which are induced against intracellular pathogens and drive proinflammatory immune effector functions aimed at pathogen clearance (29). However, T-bet expressing T cells have also been described in the context of homeostasis in the intraepithelial compartment of the intestine and mammary glands, where they participate in immune tolerance and tissue function (3, 30). To determine whether SILP CD4⁺T-bet⁺ T cells depend on classical Th1 factors, we tested their dependency on cDC1 and IL-12. Similarly to classical Th1 cells, SILP CD4⁺T-bet⁺ cells where significantly reduced in *Batf3^-/-^* mice (Figure S3A-C), which lack a transcription factor that is necessary for cDC1 cell development (31, 32). Surprisingly, the absence of IL-12p40, which is a subunit of the Th1 inducing cytokine IL-12, did not impact the accumulation of CD4⁺T-bet⁺ T cells in the small intestine (Figure S3D-E), suggesting distinct differentiation signals compared to Th1 cells.

IL-27 is a type I cytokine that is involved in both pro and anti-inflammatory functions, and which is present in the small intestine (33, 23, 22). Furthermore, a recent report showed that cDC1-derived IL-27 drives the accumulation of Th1 cells in this small intestine (34). Indeed, when we compared the CD4 T cell compartment in the small intestine between WT and cohoused *Il27ra^-/-^* mice, we observed a stark decrease in the accumulation of this population (Figure S3F-G). Finally, we tested whether T-bet itself was necessary by employing T-bet-ZsGreen-*Tbx21^-/-^* transgenic mice, which report on T-bet expression competency in the absence of T-bet expression from the endogenous locus. The frequency and number of T-bet competent CD4⁺ T cells was markedly reduced in the absence of endogenous T-bet expression (Figure S3H-I), showing that T-bet itself is required for the accumulation of this population.

These data clearly establish that homeostatic SILP CD4⁺T-bet⁺ T cells have distinct differentiation requirements compared to classical Th1 cells. Instead, this population depends on IL-27, a cytokine witch imparts antiinflammatory functions in T cells and drives the differentiation of Tr1 cells, a regulatory IL-10-secreting T cell subset that lacks FOXP3 expression (19). The requirement of IL-27 signaling for homeostatic SILP CD4⁺T-bet⁺ T cell accumulation coupled with their dependence on dietary factors raised the possibility that they may harbor unique effector functions geared towards tolerance or tissue function.

### SILP CD4⁺T-bet⁺ T cells present a suppressive signature during activation and homeostasis

To uncover the functional capacity of SILP CD4⁺T-bet⁺ T cells we performed bulk RNA sequencing of sorted CD4⁺T-bet⁺ T cells after in vivo activation (Figure 4A). Remarkably, in vivo activation led to the upregulation of a variety of suppressive genes typically associated with T regulatory cells (Figure 4B-D). Despite a strong signature of proliferation (Figure S4A), IL-10 and multiple coinhibitory receptors including *Lag3*, *Havcr2* (Tim3), *Tigit*, and *Ctla4* were among the top upregulated genes (Figure 4B-D). While we detected very few T-bet expressing Tregs at homeostasis (Figure 1A), the possibility remained that Tregs would upregulate T-bet during in vivo activation and lead to the observed results in Figure 4B-D. Therefore, we profiled SILP CD4⁺ T cells during in vivo activation by single cell RNA sequencing in both SPF and germ-free conditions (Figure 4E and S4B). This approach confirmed that upregulation of IL-10 and coinhibitory receptors occurred in T-bet⁺Foxp3⁻ cells (Figure 4G-F and S4C-D). Interestingly, we observed a similar suppressive signature in Th17 cells during in vivo activation in SPF conditions (Figure 4H-F), reminiscent of the recently published study on SFB-specific IL-10-producing cells (18). To confirm IL-10 secretion capability of SILP CD4⁺T-bet⁺ T cells, we crossed T-bet-AmCyan FoxP3-RFP reporter mice to IL-10-GFP reporter mice. In vivo T cell stimulation resulted in marked increase in the percentage of IL-10 producing SILP CD4⁺T-bet⁺ T cells, while the percentage of IL-10 producing Tregs remained unchanged (Figure 4I-K). These data argue that SILP CD4⁺T-bet⁺ T cells are in fact T-bet⁺ Tr1 cells an may harbor unique effector functions geared towards tolerance or tissue function.

**Figure 4:**
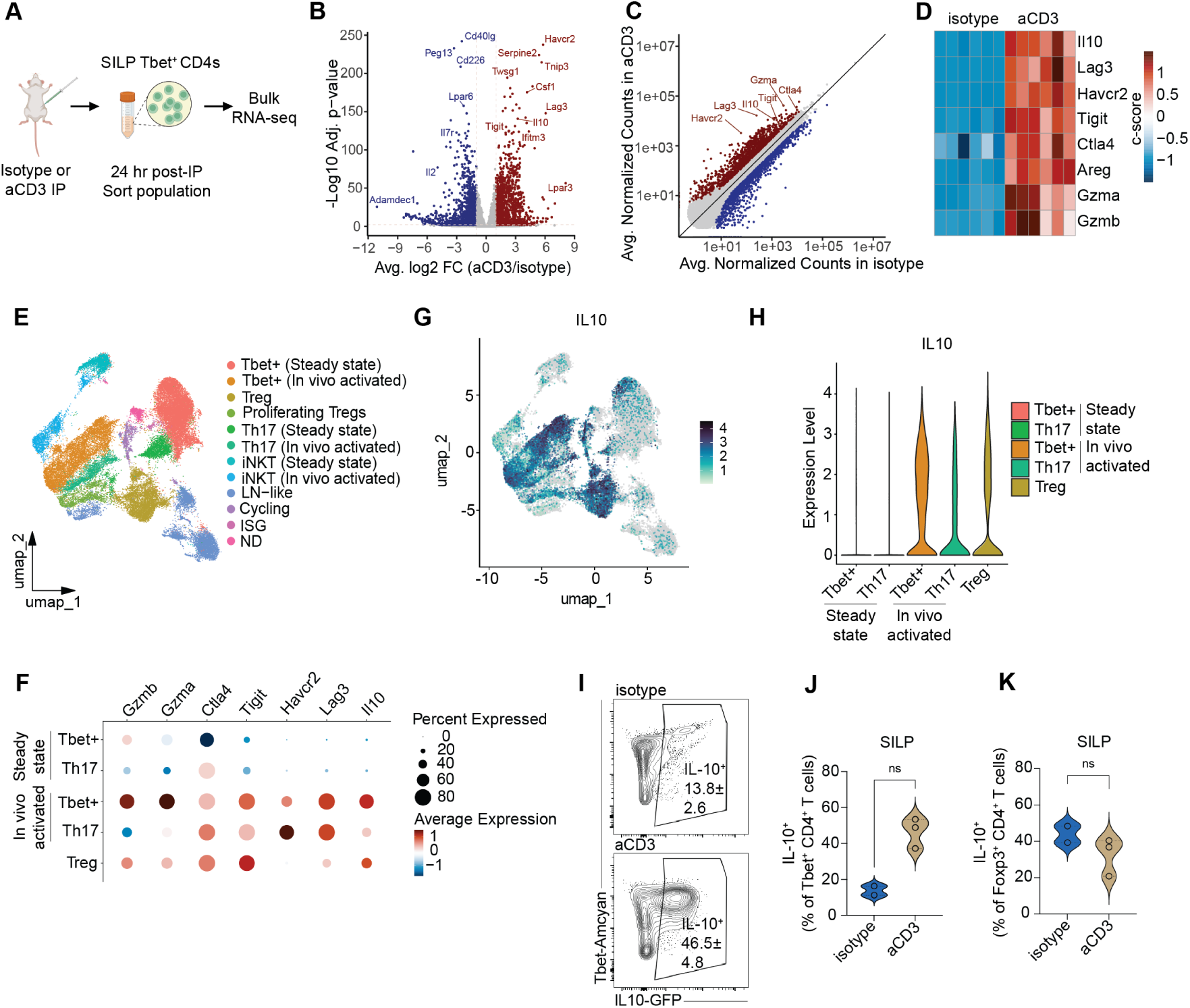
In vivo activation reveals a suppressive signature and IL-10 expression capacity of small intestinal CD4⁺T-bet⁺ T cells. (A-D) Bulk RNA sequencing of sorted SILP CD4⁺Tbet+ T cells from SPF C57BL/6-Tbet-ZsGreen mice injected with isotype or ⍺CD3 antibody 24 hours prior to tissue collection (6 mice per group). (A) Model depicting experimental design. (B) Volcano Plot showing differentially expressed genes in sorted SILP CD4⁺Tbet+ T cells from ⍺CD3- vs isotype-injected mice as measured by bulk RNAseq. (C) Scatter Plot showing differentially expressed genes in sorted SILP CD4⁺Tbet+ T cells from ⍺CD3- vs isotype-injected mice as measured by bulk RNAseq. Genes indicative of a suppressive signature are highlighted. (D) Heat map showing genes indicative of a suppressive signature in sorted SILP CD4⁺Tbet+ T cells from ⍺CD3- vs isotype-injected mice as measured by bulk RNAseq. (E-H) Single Cell RNA sequencing of sorted SILP CD4⁺ T cells from SPF and GF C57BL/6-Tbet-ZsGreen mice injected with isotype or ⍺CD3 antibody 24 hours prior to tissue collection (3-4 mice per group). (E) Uniform Manifold Approximation and Projection (UMAP) plot of SILP CD4⁺ T cells from ⍺CD3- vs isotype-injected mice. Clusters are annotated into 12 cell types. (F) Dot plot showing the expression of genes indicative of a suppressive signature in the indicated clusters. Dot size represents the faction of the cells where the gene is detected, and the color scale represents column z-scored average expression. (G) Single cell expression of *Il10* projected on the UMAP plot of SILP CD4⁺ T cells from ⍺CD3- vs isotype-injected mice. Cells where *Il10* expression is not detected are shown in gray. (H) Violin Plot showing the distribution of IL-10 expression across steady state and in-vivo activated Tbet+ and Th17; and Treg clusters. (I-K) SPF C57BL/6-T-bet-Amcyan-Foxp3-RFP-IL-10-GFP were injected with isotype or ⍺CD3 antibody 23 hours prior to tissue collection. SILP immune cells were isolated and stained with immunofluo-rescent antibodies and analyzed with flow cytometry. (I) Representative flow cytometry plots of Tbet+ Foxp3-CD4 T cells showing expression of IL-10 in isotype and ⍺CD3 groups. (J) Quantification of the frequency of IL-10-expressing CD4⁺T-bet⁺Foxp3⁻ T cells. (K) Quantification of the frequency of IL-10-expressing CD4⁺Foxp3⁺ T cells (Tregs). Data are representative of at least three independent experiments (I-K). Ns, not significant, *P < 0.0332, **P < 0.0021, ***P < 0.0002, ****P < 0.0001. Comparisons used Mann-Whitney test. Error bars indicate SEM.

Interestingly, even at homeostasis a small but detectable proportion of the highest T-bet expressing CD4⁺ T cells produced IL-10 (Figure 4I). Given the role the dietary components played in the accumulation and clonal selection of this population (Figure 3), we hypothesized that diet may also drive the suppressive program and homeostatic IL-10 secretion observed in SILP Tr1 cells. Indeed, comparing their transcriptional profile between mice raised on a regular chow, casein, or amino acid diet by pseudobulk analysis of single cell RNA sequencing data revealed an upregulation of the suppressive signature in the regular chow condition (Figure S4E-J). Collectively, our results uncover a suppressive potential of T-bet expressing CD4⁺ T cells in the small intestine characterized by the secretion of IL-10 and driven by dietary signals, unveiling this population as T-bet⁺ Tr1 cells.

## Discussion

In this work we describe a previously unappreciated T-bet expressing Tr1 T cell population in the small intestine which represents an outsized arm of homeostatic T cell immunity in this tissue. Of particular importance, this population is entirely independent of the microbiota, and is instead controlled by dietary signals. This stands in stark contrast to other homeostatic T cell responses described to date in the small and large intestines (1, 2), where particular commensal species drive cognate effector and regulatory T cell immunity. Instead, T-bet⁺ Tr1 cells mirror small intestinal Tregs in their dependence on diet and IL-10 expression, raising the possibility that immune tolerance in the small intestine is not sole purvey of T regulatory cells and instead encompasses Foxp3-negative T-bet-expressing Tr1 cells, with important potential implications for the understanding of immunoregulation and oral tolerance.

Early life represents a critical window for the education and development of the immune system (26), (25). Mounting evidence demonstrates that early life environmental and microbial exposures imprint long-term immunological set points that shape immune function and disease susceptibility into adulthood (24). In the intestine, the transition from maternal milk to solid food induces a rapid remodeling of intestinal tissue physiology, immune function, and microbiota composition (27). Indeed, antigen uptake and presentation in the small and large intestines are carefully regulated during this time period, contributing to tolerance induction towards commensal and environmental antigens (35). Concomitantly, there is an acute influx of T lymphocytes into the intestine (28) as part of the establishment of long-term homeostatic immune function in this tissue. Here we reveal that T-bet expressing T cells dominate the small intestinal immune landscape at homeostasis, and we describe T-bet⁺ Tr1 cells as a major component of small intestinal T cell immunity established upon weaning.

The microbiota plays a critical role in establishing the T cell immune compartment in the intestine (1). However, we show that T-bet⁺ Tr1 cells are not controlled by microbial signals, as their accumulation, TCR repertoire, and function remains unchanged in germ-free conditions. Instead, dietary components drive their accumulation, clonal selection, and suppressive program in the small intestine. T-bet⁺ Tr1 cells mirror small intestinal regulatory cells, which accumulate in a spimilar timeline in response to dietary antigens and participate in oral tolerance to food antigens. Importantly, these cells are enriched proximally in the duodenum, a site for digestion and absorption of dietary nutrients which harbors a very small bacterial load, consistent with their dependence on dietary antigens and their independence of the microbiota. Whether T-bet⁺ Tr1 cells recognize dietary polypeptide antigens directly, or whether they may be specific for self or a environmental antigens and respond to dietary derived signals will be important to address.

Tr1 cells are characterized by the secretion of IL-10 in the absence of Foxp3 expression and develop from effector T cells in the context of chronic antigen stimulation (19). The intestine is constantly exposed to dietary antigens, raising the possibility that the Tr1 program allows T-bet⁺ Tr1 cells to adopt an immunoregulatory phenotype, contributing to immunoregulation and/or oral tolerance. Indeed, we observed IL-10 expression by a subset of T-bet⁺ Tr1 cells at homeostasis, accompanied by a marked competency for overt IL-10 expression by most of the remaining cells in this compartment upon activation. IL-10 expression was accompanied by upregulation of inhibitory costimulatory receptors such as Lag3, Tim3, Ctla4, and Tigit, suggesting that suppressive functions of T-bet⁺ Tr1 cells may extend beyond IL-10 secretion. Indeed, a previous report identified the expression of inhibitory receptors by IL-10 producing Tr1 cells as a predictive marker of suppressive capacity(36). Still, a subset of T-bet⁺ Tr1 cells had the capacity for IFN𝛾 and granzyme expression at steady state. Whether more classical T-bet regulated functions are provided by T-bet⁺ Tr1 cells for other purposes remains an open question. Still, the suppressive signature was dominant upon activation, and extended beyond T-bet⁺ Tr1 cells to Th17 cells, as described in the literature for SFB-specific responses (18). Together, these observations support the hypothesis that an immunoregulatory program may be a shared feature of small intestinal T cell immunity, perhaps driven by local cues.

The Tr1 program is driven by a combination of transcription factors and upstream signals, including c-Maf, Ahr, and IL-27 (20). Ahr ligands are abundant in the small intestine (21), and we detected Ahr and c-Maf expression across small intestinal CD4⁺ T cells. IL-27 is abundant in the small intestine [J. Diegelmann, T. Olszak, B. Göke, R. S. Blumberg, S. Brand. (37)](22, 23), and a recent report revealed that cDC1 derived IL-27 controls Th1 accumulation in the intestine (34) at homeostasis. While IL-27 can promote proinflammatory functions in certain contexts, it remained unclear why classical Th1 cells in the small intestine would depend on this cytokine. Our work reveals that IL-27 driven CD4⁺T-bet⁺ T cells are not classical Th1 cells, but are in fact IL-10 secreting T-bet⁺ Tr1 cells. Furthermore, in contrast to antibiotic mediated approaches employed in that report (34), our data conclusively show that this population does not depend on the microbiota. Differences in our results regarding this matter may be due to off target effects of antibiotic treatment. Alternatively, a fraction of CD4⁺T-bet⁺ T cells at homeostasis could be driven by facility-restricted commensals or pathogens, which would be depleted by antibiotic treatment. Notwithstanding these observations, our data conclusively show that the majority of CD4⁺T-bet⁺ T cells in the small intestine are not controlled by the microbiota. Furthermore, our work identifies T-bet not just as a unique marker of this population but as a critical regulator of the accumulation in the small intestine. One possibility is that T-bet controls the expression of certain chemokine receptors or integrins which mediate recruitment into the small intestine. Furthermore, a recent study examining Tr1 transcriptional networks in vitro identified T-bet as a candidate regulator of the Tr1 program (20), which would agree with our in vivo observations and place T-bet at the center of the transcriptional program of SILP Tr1 cells.

Immune regulation in the intestine is critical to safeguard tissue function. Oral tolerance of dietary antigens represents a critical function of intestinal immunity, involving the induction of T regulatory responses to dietary antigens. Breakdown of oral tolerance leads to life threatening food allergy or chronic intestinal autoimmunity such as celiac disease (5). The mechanisms governing the induction and maintenance of oral tolerance both systemically and in the intestine remain an active area of investigation, and have largely focused on T regulatory cells (7, 6). Our work raises the possibility that other T cell subsets beyond Tregs, such as small intestinal T-bet⁺ Tr1 cells may participate in immune regulation in the small intestine, potentially extending to oral tolerance. Thus, understanding the antigen specificity and effector functions of SILP T-bet⁺ Tr1 cells may provide critical insights into the mechanisms of immune regulation, autoimmunity, oral tolerance, and food allergy, paving the way for innovative interventions and therapies that harness this T cell population.

## Acknowledgments

This work was supported by the NIAID Division of Intramural Research. We thank the NIAID animal facility staff for animal husbandry; G. Koroleva, T. Hawley, and J. Polanco (Center for Human Immunology) for technical assistance; and Eric Dang (NIAID) for providing feedback on the manuscript. We thank the NIH tetramer core facility for providing tetramers. We thank all members of the Belkaid laboratory for helpful scientific discussions.

## Data availability

All sequencing date have been deposited to GEO, with the following accession numbers: - GSE298371: Single-cell RNA sequencing of SILP CD4 T cells from SPF and GF mice. - GSE298394: Bulk RNA Sequencing of invivo activated SILP CD4⁺T-bet⁺ T cells. - GSE298386: Single-cell TCR sequencing of SILP CD4⁺T-bet⁺ T cells from GF mice raised on casein, amino acid, or regular chow diets. - GSE298374: Single-cell RNA sequencing of Small Intestinal Lamina Propria invivo activated CD4 T cells from SPF and GF mice.

## Materials and Methods

### Mice

Wild-type C57BL/6, CD45.1 (Taconic line 8478), T-bet-Zsgreen (Taconic Line 8419), I-Ab-/-, BATF3-/-, Il-12p40-/-, Il27ra-/-, and Tbx21-/- mice were purchased from Taconic and maintained at NIAID animal facilities. All mice were bred and maintained under specific pathogen-free or germ-free conditions at an American Association for the Accreditation of Laboratory Animal Care (AAALAC)-accredited animal facility at NIAID and housed in accordance with the procedures outlined in the Guide for the Care and Use of Laboratory Animals. All experiments were performed at NIAID under an animal study proposal (LHIM-2E) approved by the NIAID Animal Care and Use Committee. Mice were group housed (4–5 mice of same sex per cage). Mice were randomly assigned to each experimental group. Unless otherwise noted, sex- and age-matched mice between 8 and 12 weeks of age were used for each experiment. Both male and female mice were included in experimental groups.

### Timed pregnancy model

Male mice were single housed for 1-2 weeks to accumulate dirty bedding and sperm recuperation. On day 0, female mice were intraperitoneally injected with 2IU of PSMG and transferred into an empty dirty male cage. 48 hours after the injection, females were intraperitoneally injected with 2IU of hCG and housed with one single male for overnight mating. The next day, females were returned to their previous cages. Female mice were checked for pregnancy 12 days after mating.

### Tissue preparation and cell extraction

Serum-free media was prepared with RPMI containing 20 mM HEPES, L-Glutamine, 100U/mL Penicillin, 100ug/mL Streptomycin sulfate, 50 µM β-mercaptoethanol, 1mM sodium pyruvate, 1mM non-essential amino acids.3% media was prepared with RPMI containing 3% FBS, 20 mM HEPES, L-Glutamine, 100U/mL Penicillin, 100ug/mL Streptomycin sulfate, 50 µM β-mercaptoethanol, 1mM sodium pyruvate, 1mM non-essential amino acids. Mice were sacrificed by CO2 inhalation. For cell extraction from small intestinal lamina propria, Peyer’s patches and mesenteric adipose tissue were removed. For each intestine, tissues were cut open longitudinally, washed several times in cold PBS to remove feces, and further cut into 1- to 2-cm segments before being placed in 50 ml beakers with a stir bar. Tissue pieces were incubated in 10 ml per intestine of 3% RPMI with 5 mM EDTA, and 0.145 mg/ml DTT for 20 min at 37° C with stirring ∼400 rpm. To separate the lamina propria (LP) and intestinal epithelial lymphocyte (IEL) layer, each beaker’s contents were strained through a sterile fine-meshed stainless-steel sieve into a corresponding 500 ml beaker containing 10 ml of 3% RPMI media held on ice. After straining, tissue pieces from each intestine were transferred to a 50 ml conical tube containing 10 ml of serum-free RPMI with 2 mM EDTA. Each tube was shaken vigorously for 30 seconds and then strained through the respective sieve into the corresponding beaker. The shake and strain processes were repeated two more times. To isolate the IEL compartment, strained contents were filtered through 70 µm cell strainers atop 50 ml conical tubes on ice. Cell suspensions were centrifuged for 5 minutes at 450 g, 4° C. After gently discarding the supernatant, cell pellets were resuspended in 4 ml of a 30% Percoll gradient and centrifuged for 5 minutes at 1800 rpm, 4° C. After removing the supernatant with a pipette, cells were resuspended in 1 ml of 3% RPMI media. To isolate the LP compartment, small intestine pieces recovered from the shake and strain steps were placed into a 50 ml beaker and finely minced with scissors. Tissues were then incubated with 10 ml of serum-free media with 0.1 mg/ml Liberase TL and 5 mg/ml DNAse I for 25 min at 37° C with stirring ∼400 rpm. After the incubation, 5 ml of 3% RPMI was added to each beaker. The contents of each beaker were pipetted through 70 µm cell strainers atop 50 ml conical tubes on ice. Each beaker and strainer was rinsed with an additional 10 ml of 3% RPMI. Cell suspensions were centrifuged for 5 minutes at 450 g, 4° C. After gently discarding the supernatant, cell pellets were resuspended in 4 ml of a 30% Percoll gradient and centrifuged for 5 minutes at 1800 rpm, 4° C. After removing the supernatant with a pipette, cells were resuspended in 1 ml of 3% RPMI media.

### Flow cytometry

Single cell suspensions were incubated with fluorochrome-conjugated antibodies against surface markers: B220 (RA3-6B2), CD4 (RM4-5), CD8a (53-6.7), CD8b (H35-17.2), CD11b (M1/70), CD11c (N418), CD44 (IM7), CD45 (30-F11), CD90.2 (30-H12), Ki-67 (SolA15), TCRb (H57-597), and TCRgd (GL3) in PBS containing 2% FBS, 1mM EDTA for 30 min at 4° C and then washed. For intracellular staining, cells were fixed and permeabilized with the Foxp3/Transcription Factor Staining Buffer Set (eBioscience) and stained with fluorochrome-conjugated antibodies: Foxp3 (FHJ-16s), GATA3 (TWAJ), Granzyme B (QA16A02), IFNγ (XMG1.2), RORγt (Q31-378), IL-17A (17B7), T-bet (4B10), and TNF (MP6-XT22) for 45 min at RT and then washed. All staining were performed in the presence of purified (fc block) anti-mouse CD16/32 (2.4G2, BioXcel). Dead cells were excluded from live samples using 4’,6-diamidino-3-phenylindol (DAPI; Sigma) whereas a LIVE/DEAD Fixable Blue Dead Cell Stain Kit (Invitrogen Life Technologies) was used in fixed samples. All antibodies were purchased from BD Biosciences, Biolegend, eBioscience or invitrogen. Flow cytometric data were acquired on a BD LSRFortessa and analyzed using FlowJo software (BD). Cell sorting was performed on a Sony MA900 sorter.

### Special diets

All mice had free access to a pelleted rodent diet and water. Breeders were set up on an amino acid, casein, or control (standard chow) diet ad libitum. Pups from these breeders were weaned onto the same corresponding diets. At 8-12 weeks of age, mice were euthanized, and the small intestine harvested and processed. The primary diet used for our control diet, we used the Global 16% Protein Rodent Diet manufactured by Inotiv Teklad (TD.00217), containing protein derived from complex proteins. For the amino acid diet, we used a Casein Mimic AA 93G Diet manufactured by Inotiv Teklad, (TD.210529) (Supplementary table 1), containing a protein source restricted to free amino acids. For the casein diet, we used an AIN-93G Diet manufactured by Inotiv Teklad (TD.210528) (Supplementary table 2), containing a refined protein source of casein.

### In-vivo activation

For experiments analyzing in-vivo activated T cells, mice were injected intra-peritoneally with 50 ug of aCD3 or isotype antibody, 23 hours prior to tissue harvesting. In-vivo activation performed using InVivoPlus 145-2C11 anti-mouse aCD3ε monoclonal antibody manufactured by Bio X Cell. The isotype control using InVivoPlus polyclonal Armenian hamster IgG isolated from Armenian hamster serum, manufactured by Bio X Cell. Each mouse was injected with 50ug of antibody in 100uL sterile 1X Phosphate-Buffered Saline (PBS).

### Single-cell RNA sequencing of Small Intestinal Lamina Propria CD4 T cells from SPF and GF mice (Related to Figure 2)

SILP CD4 T cells (Live, CD45+CD90.2+, TCRb+, CD4+) from 4 SPF and 4 GF T-bet-ZsGreen mice were sorted from samples labeled with TotalSeqC hashtags antibodies (Biolegend) using a Sony MA900 sorter. Sorted cells were pooled during sorting and 32000 cells per lane (total of 8 lanes) were loaded on a Chromium Single Cell Controller (10X Genomics) to encapsulate the cells into droplets. The gene expression (GEX), hashtag oligonucleotide (HTO) and VDJ (TCR) sequencing libraries were prepared using Chromium Single Cell 5’ v2 reagent kits following the manufacturer’s instructions, and sequenced on an Illumina Nextseq2000 platform (NextSeq 1000/2000 P2 reagents). The initial processing of the expression data involved generating fastq files and count matrices using Cell Ranger v7.1.0 (38) (10X Genomics, Pleasanton, CA) that was run with the “Include introns=True” option and the mm10-2020-A transcriptome reference. The TCR data was processed with Cell Ranger v7.1.0 using a VDJ reference based on the IGMT database: The IGMT reference was generated with Cell Ranger v7.1.0 using the ‘fetch-imgt’ and ‘cellranger mkvdjref’ functions. Processing of the expression data was performed with the ‘cellranger multi’ and ‘cell-ranger aggr’ functions. After processing and aggregation, the estimated number of cells and mean reads per cell for each library were as follows: 89077 estimated number of cells, 19550 mean reads per cell (from eight GEX libraries), 1208 Median hashtag UMIs per Cell (across eight HTO libraries), 56835 lls with a productive TRA and/or TRB V-J spanning sequence, 38232 of which contain both a productive V-J spanning TRA and TRB pair (from eight VDJ libraries). The overall sequencing quality was medium to high in the eight mRNA libraries: >82.7% of bases in the barcode and UMI regions had Q30 (99.99% inferred base call accuracy, mean 94%) quality score or above, whereas >75.4% of bases in the RNA reads had Q30 or above (mean 85.6%). Furthermore, across the eight GEX libraries the median gene count per cell range was 974-1332, the sequencing saturation ranged between 69-76%, and 39-54% of the reads mapped confidently to the transcriptome. Across the eight TCR libraries, the mean used reads per cell was 1771-4034 (average 2755). Across the eight hashtag (HTO) libraries the mean antibody reads usable per cell was 1956-6321 (average 3015).

The downstream analysis of the expression data was performed in R 4.4.2, the Cell Ranger aggregation output (filtered_feature_bc_matrix directory) was loaded into Seurat version 5.1.0 (39) using the CreateSeuratObject function with min.cells=3. For filtering out low-quality cells, we established maximum thresholds for gene and UMI counts (nCount_RNA < 25000, 200 < nFeature_RNA < 5000), and a maximum threshold for percentage mitochondrial content (percent.mt < 6) based on visual inspection of the corresponding metric distributions across all cells. Log-normalization (LogNormalize) of the RNA count data and centered log-ratio transformation (CLR) of the HTO count data were carried out using the NormalizeData function. For HTO demultiplexing cells into the individual mice within SPF and GF groups and removing any doublets, HTODemux was run with a positive quantile = 0.99 and only cells identified as Singlets were kept. The top 2000 variable features were determined with the VariableFeatures function, expression data was scaled with ScaleData, and dimensionality reduction was performed with the RunPCA function on the variable features, followed by determining the dimensionality of the dataset on an elbow plot (ElbowPlot function), which was determined to be 75. Subsequently, FindNeighbors, FindClusters, and RunUMAP were run on 75 dimensions and a range of resolutions to determine the optimal resolution that captures biological diversity in the dataset, which was determined to be 0.25, resulting in 20 clusters. The clusters identified by the FindClusters function of Seurat (default Louvain clustering setting and 0.25 resolution) were annotated using cell type specific markers, cluster specific differentially expressed genes, and with the aid of SingleR (40), using the ImmGen database reference. Contaminating clusters comprised of B cell, endothelial, CD8+ T cells, Macrophages, and stromal fibroblast cells were removed. On the second iteration of clustering andremoval of contaminating cell types, small clusters corresponding to ILCs and low quality cells that were possible doublets with B cells were removed. On the remaining cells FindVariableFeatures (nFeatures=2000) was run, data was scaled with ScaleData, RunPCA was run (npcs=100) and the top 38 dimensions were used to run FindNeighbors, FindClusters (with a range of resolutions and a final chosen resolution of 0.25), and RunUMAP. The resulting clusters were annotated using cell type specific markers, cluster specific differentially expressed genes, and with the aid of SingleR (40), using the ImmGen database reference

### Single-cell TCR sequencing of Small Intestinal Lamina Propria CD4⁺T-bet⁺ T cells from GF mice raised on casein, amino acid, or regular chow diets (Related to Figure 3)

SILP CD4⁺T-bet⁺ T cells (Live, CD45+CD90.2+, TCRb+, CD4+, T-bet-ZsGreen+) from GF T-bet-ZsGreen mice raised on casein, amino acid, or regular chow diet were sorted from samples labeled with TotalSeqC hashtags antibodies (Biolegend) using a Sony MA900 sorter. Sorted cells were pooled during sorting and 40000 cells per lane (total of 4 lanes) were loaded on a Chromium Single Cell Controller (10X Genomics) to encapsulate the cells into droplets. The gene expression (GEX), hashtag oligonucleotide (HTO) and VDJ (TCR) sequencing libraries were prepared using Chromium Single Cell 5’ v2 reagent kits following the manufacturer’s instructions, and sequenced on an Illumina Nextseq1000 platform (NextSeq 1000/2000 P2 reagents). The initial processing of the expression data involved generating fastq files and count matrices using Cell Ranger v7.1.0 (38) (10X Genomics, Pleasanton, CA) that was run with the “Include introns=True” option and the mm10-2020-A transcriptome reference. The TCR data was processed with Cell Ranger v7.1.0 using a VDJ reference based on the IGMT database: The IGMT reference was generated with Cell Ranger v7.1.0 using the ‘fetch-imgt’ and ‘cellranger mkvdjref’ functions. Processing of the expression data was performed with the ‘cellranger multi’ and ‘cellranger aggr’ functions. After processing and aggregation, the estimated number of cells and mean reads per cell for each library were as follows: 56451 estimated number of cells, 24386 mean reads per cell (from 4 GEX libraries), 609 Median hashtag UMIs per Cell (across 4 HTO libraries), 43091 cells with a productive TRA and/ or TRB V-J spanning sequence, 34070 of which contain both a productive V-J spanning TRA and TRB pair (from 4 VDJ libraries). The overall sequencing quality was high in the 4 mRNA libraries: >95% of bases in the barcode and UMI regions had Q30 (99.99% inferred base call accuracy) quality score or above, whereas >94% of bases in the RNA reads had Q30 or above. Furthermore, across the 4 GEX libraries the median gene count per cell range was 1835-2018, the sequencing saturation ranged between 68-78%, and 73-76% of the reads mapped confidently to the transcriptome. Across the 4 TCR libraries, the mean used reads per cell was 1107-5279 (average 3201). Across the 4 hashtag (HTO) libraries the mean antibody reads usable per cell was 1165-1486 (average 1319).

The downstream analysis of the expression data was performed in R 4.4.2, the Cell Ranger aggregation output (filtered_feature_bc_matrix directory) was loaded into Seurat version 5.1.0 (39) using the CreateSeuratObject function with min.cells=3. For filtering out low-quality cells, we established maximum thresholds for gene and UMI counts (nCount_RNA > 100, nFeature_RNA > 500), and a maximum threshold for percentage mitochondrial content (percent.mt < 5) based on visual inspection of the corresponding metric distributions across all cells. Log-normalization (LogNormalize) of the RNA count data and centered log-ratio transformation (CLR) of the HTO count data were carried out using the NormalizeData function. For HTO demultiplexing cells into individual mice and removing any doublets, HTODemux was run with a positive quantile = 0.99 and only cells identified as Singlets were kept. The top 2000 variable features were determined with the VariableFeatures function, expression data was scaled with ScaleData, and dimensionality reduction was performed with the RunPCA function on the variable features, followed by determining the dimensionality of the dataset on an elbow plot (ElbowPlot function), which was determined to be 70. Subsequently, FindNeighbors, FindClusters, and RunUMAP were run on 70 dimensions and a range of resolutions to determine the optimal resolution that captures biological diversity in the dataset, which was determined to be 0.5, resulting in 26 clusters. The clusters identified by the FindClusters function of Seurat (default Louvain clustering setting and 0.5 resolution) were annotated using cell type specific markers, cluster specific differentially expressed genes, and with the aid of SingleR (40), using the ImmGen database reference. Contaminating clusters comprised of B cells and epithelial cells were removed. On the second iteration of clustering and removal of contaminating cell types, Tregs were removed so that only CD4⁺T-bet⁺Foxp3⁻ T cells remained. On the remaining cells FindVariableFeatures (nFeatures=2000) was run, data was scaled with ScaleData, RunPCA was run (npcs=100) and the top 45 dimensions were used to run FindNeighbors, FindClusters (with a range of resolutions and a final chosen resolution of 0.15), and RunUMAP.

Pseudobulk analysis on CD4⁺T-bet⁺Foxp3⁻ T cells with the indicated comparisons was performed as described in the corresponding methods with a FC threshold of 0.3 and an adjusted p value threshold of 0.05.

### Single-cell RNA sequencing of Small Intestinal Lamina Propria invivo activated CD4 T cells from SPF and GF mice (Related to Figure 4)

SILP CD4⁺ T cells (Live, CD45+CD90.2+, TCRb+, CD4+) from SPF or GF T-bet-ZsGreen mice that received aCD3 or isotype as described in the corresponding methods section were sorted from samples labeled with TotalSeqC hashtags antibodies (Biolegend) using a Sony MA900 sorter. Sorted cells were pooled during sorting and 40000 cells per lane (total of 4 lanes) were loaded on a Chromium Single Cell Controller (10X Genomics) to encapsulate the cells into droplets. The gene expression (GEX), hashtag oligonucleotide (HTO) and VDJ (TCR) sequencing libraries were prepared using Chromium Single Cell 5’ v2 reagent kits following the manufacturer’s instructions, and sequenced on an Illumina Nextseq1000 platform (NextSeq 1000/2000 P2 reagents). The initial processing of the expression data involved generating fastq files and count matrices using Cell Ranger v7.1.0 (38) (10X Genomics, Pleasanton, CA) that was run with the “Include introns=True” option and the mm10-2020-A transcriptome reference. The TCR data was processed with Cell Ranger v7.1.0 using a VDJ reference based on the IGMT database: The IGMT reference was generated with Cell Ranger v7.1.0 using the ‘fetch-imgt’ and ‘cellranger mkvdjref’ functions. Processing of the expression data was performed with the ‘cellranger multi’ and ‘cellranger aggr’ functions. After processing and aggregation, the estimated number of cells and mean reads per cell for each library were as follows: 74799 estimated number of cells, 20476 mean reads per cell (from 4 GEX libraries), 1261 Median hashtag UMIs per Cell (across 4 HTO libraries), 48717 cells with a productive TRA and/or TRB V-J spanning sequence, 38824 of which contain both a productive V-J spanning TRA and TRB pair (from 4 VDJ libraries). The overall sequencing quality was high in the 4 mRNA libraries: 68-97% of bases in the barcode and UMI regions had Q30 (99.99% inferred base call accuracy, average 92%) quality score or above, whereas 68-95% (average 90) of bases in the RNA reads had Q30 or above. Furthermore, across the 4 GEX libraries the median gene count per cell range was 1774-1963, the sequencing saturation ranged between 48-53%, and 76-79% of the reads mapped confidently to the transcriptome. Across the 4 TCR libraries, the mean used reads per cell was 1306-4043 (average 2112). Across the 4 hashtag (HTO) libraries the mean antibody reads usable per cell was 2532-3912 (average 2960).

The downstream analysis of the expression data was performed in R 4.4.2, the Cell Ranger aggregation output (filtered_feature_bc_matrix directory) was loaded into Seurat version 5.1.0 (39) using the CreateSeuratObject function with min.cells=3. For filtering out low-quality cells, we established maximum thresholds for gene and UMI counts (nCount_RNA > 1200, nFeature_RNA > 500), and a maximum threshold for percentage mitochondrial content (percent.mt < 5) based on visual inspection of the corresponding metric distributions across all cells. Log-normalization (LogNormalize) of the RNA count data and centered log-ratio transformation (CLR) of the HTO count data were carried out using the NormalizeData function. For HTO demultiplexing cells into individual mice and removing any doublets, HTODemux was run with a positive quantile = 0.99 and only cells identified as Singlets were kept. The top 2000 variable features were determined with the VariableFeatures function, expression data was scaled with ScaleData, and dimensionality reduction was performed with the RunPCA function on the variable features, followed by determining the dimensionality of the dataset on an elbow plot (ElbowPlot function), which was determined to be 45. Subsequently, FindNeighbors, FindClusters, and RunUMAP were run on 45 dimensions and a range of resolutions to determine the optimal resolution that captures biological diversity in the dataset, which was determined to be 0.25, resulting in 26 clusters. The clusters identified by the FindClusters function of Seurat (default Louvain clustering setting and 0.25 resolution) were annotated using cell type specific markers, cluster specific differentially expressed genes, and with the aid of SingleR (40), using the ImmGen database reference. Contaminating clusters comprised of B cells, fibroblasts, and epithelial cells were removed. On the remaining cells FindVariableFeatures (nFeatures=2000) was run, data was scaled with ScaleData, RunPCA was run (npcs=100) and the top 40 dimensions were used to run FindNeighbors, FindClusters (with a range of resolutions and a final chosen resolution of 0.4), and RunUMAP. The resulting 19 clusters were annotated using cell type specific markers, cluster specific differentially expressed genes, and with the aid of SingleR (40), using the ImmGen database reference.

Pseudobulk analysis on CD4⁺T-bet⁺Foxp3⁻ T cells with the indicated comparisons was performed as described in the corresponding methods section with a log2FC threshold of 1 and an adjusted p value threshold of 0.01.

### Single Cell TCR repertoire analysis

TCR repertoire sequencing data were analyzed with the scRepertoire package v2.2.1 (41) in R 4.4.2. Starting with the filtered contig annotations output table from ‘cellranger aggr’, the createHTOContigList, combineTCR, and combineExpression functions were used for combining the TCR data from each sample and for integration of the combined TCR data with the corresponding scRNAseq Seurat object. We utilized the amino acid sequence of the paired CDR3 regions of the alpha and beta chains (or a single alpha or beta chain if the other was not detected) as the clonotype definition. T cell repertoire diversity was estimated using the D50 metric, which was calculated in R as the fraction of clonotypes, ordered by abundance, which account for 50 percent of total TCR sequences. Clonotype size was defined at the level of each biological replicate (Of all sequences: Rare: between 0% and 0.01%, Small: between 0.01% and 0.1%, Medium: between 0.1% and 1%, Large: between 1% and 10%, Hyperexpanded: between 10% and 100%). Cells with a putative iNKT TCR were defined as containing a TRAV11-TRAJ18 alpha chain V gene usage, while cells with a putative MAIT TCR were defined as containing a TRAV1-TRAJ33 alpha chain V gene usage. TCR overlap between mice was defined with the same clonotype definition: at level of the CDR3 amino acid region for the combined alpha and beta chains, or a single alpha or beta chain if the other was not detected; computed in R, and visualized using the package circlize (42).

### Pseudobulk analysis of single cell RNA sequencing data

For pseudobulk analysis, the raw counts were aggregated across cells for each biological replicate on the indicated clusters using AggregateExpression using Seurat 5.1.0 (39) in R 4.4.2. Genes with a very low level of expression across biological replicates, less than 10 total counts across all samples, were filtered from the dataset. Differential gene expression analysis was performed using DESeq2 v1.44.0 (43) on the aggregated raw counts, using default parameters, and not including a log2 fold change shrinkage step. The normalized counts were extracted from the DESeq2 object and used for visualization along be corresponding adjusted p-values and log2 fold changes. Pathway enrichment analysis of upregulated or downregulated genes was performed using Metascape (44).

### Bulk RNA Sequencing of invivo activated SILP CD4⁺T-bet⁺ T cells

SILP CD4⁺ T cells (Live, CD45+CD90.2+, TCRb+, CD4+) from SPF or GF T-bet-ZsGreen mice that received aCD3 or isotype as described in the corresponding methods section were sorted using a Sony MA900 sorter. RNA was extracted using the Qiagen miRNeasy kit, and sequencing library was prepared using Tecan SoLo total RNA kit for mouse according to manufacturer’s instructions. Libraries were sequenced as 1 x 150bp reads on a Illumina NextSeq 2000 using the P3 200 cycle kit.*

Sequencing reads were mapped to the C57BL/6 mouse genome (GRCm38: mm10) and differential gene expression was calculated utilizing HOMER’s (45) getDifferentialExpression with default parameters. Differentially expressed genes (FDR < 0.01, log2FC > 1) were used for gene ontology (GO) enrichment analysis with MetaScape (44).

---

**Figure S1, related to Figure 1:**
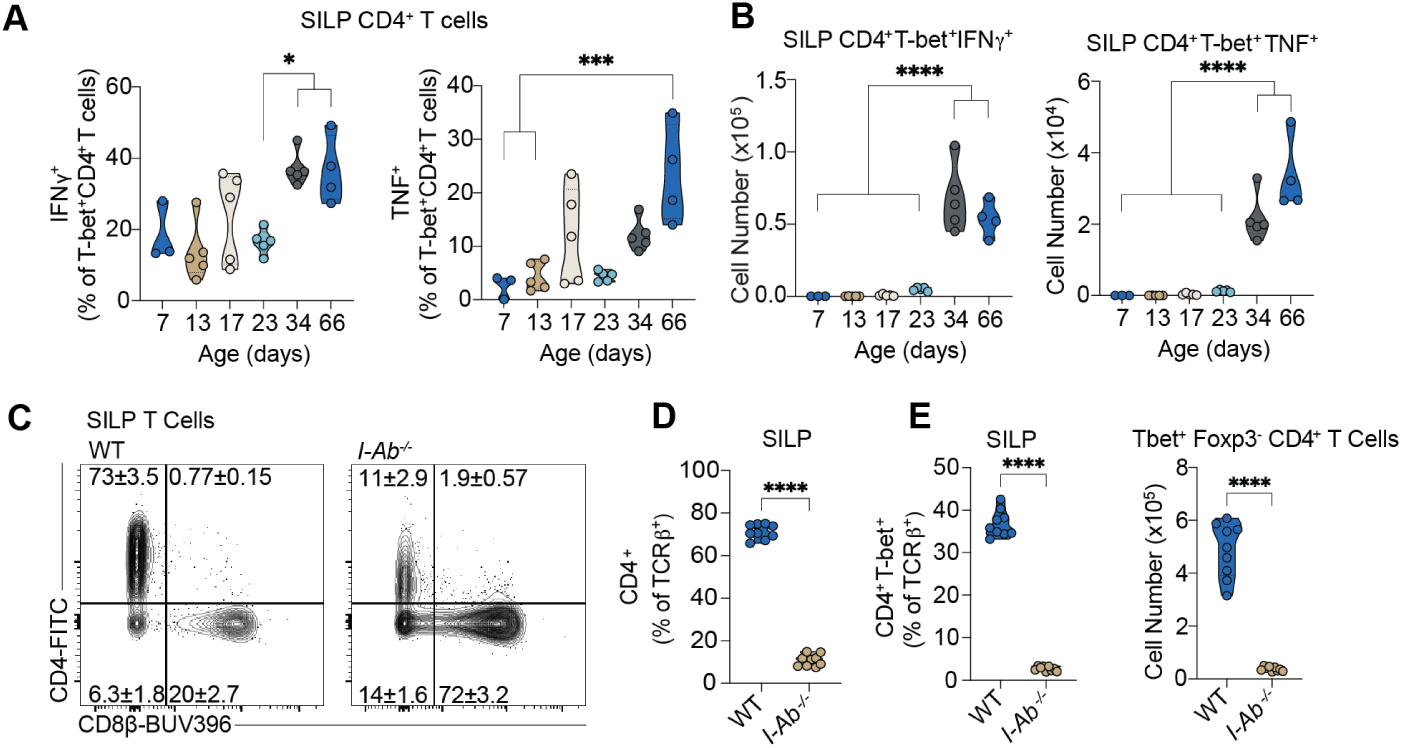
The small intestinal immune landscape is dominated by CD4⁺T-bet⁺ T cells. (A - B) Time course of IFN𝛾 and TNF production by ex vivo stimulated SILP cells throughout weaning. (A) Frequencies and (B) cell counts of cytokine production by SILP CD4+T-bet+Foxp3- T cells. (C) Representative flow plots showing CD4+ and CD8+ frequency in wildtype or *I-Ab^-/-^* SPF mice gated on CD45+ CD90.2+ TCRb+ Cd1d-T cells. (D) Violin plot showing SILP CD4+ T cell frequencies across groups for data shown in (C). (E) Violin plots showing CD4+ T-bet+ frequency and counts across wildtype or *I-Ab^-/-^* SPF mice gated on CD45+ CD90.2+ TCRb+ Cd1d-CD4+ T cells. Data are representative of at least three independent experiments. Ns, not significant, *P < 0.0332, **P < 0.0021, ***P < 0.0002, ****P < 0.0001. Brown-Forsythe and Welch ANOVA tests (A - B), unpaired t test with Welch’s correction (D - E). Error bars indicate SEM.

**Figure S2: related to Figure 2:**
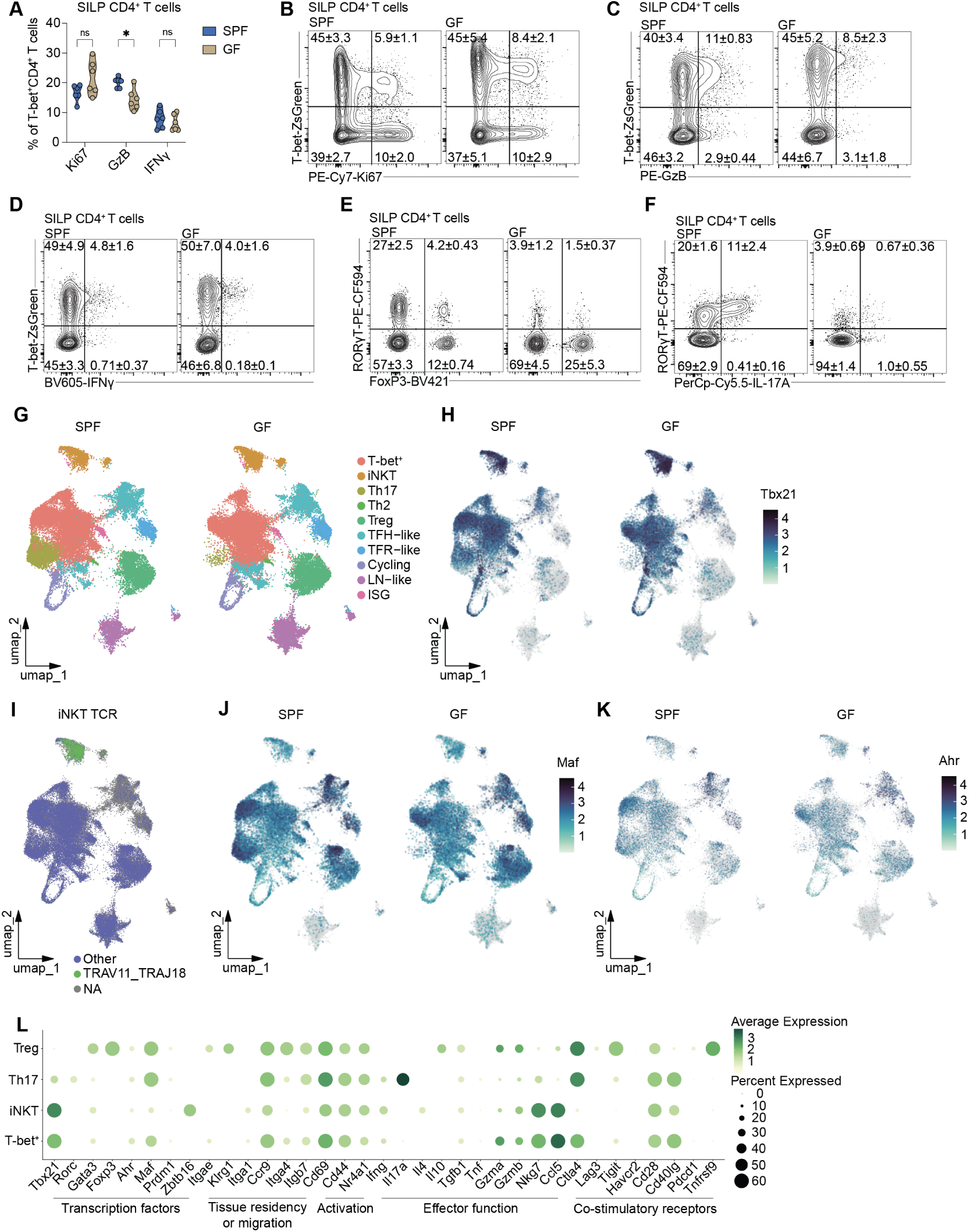
Small intestinal CD4⁺T-bet⁺ T cells are independent of the microbiota. (A) Violin plots showing frequency of Ki67-, GzB-, or IFN𝛾-expressing cells by unstimulated (Ki67, GzB) or ex vivo stimulated (IFN𝛾) SILP CD45+ CD90.2+ TCRb+ T-bet+ FoxP3-CD4+ T * cells from SPF or GF mice. (B-D) Representative flow plots showing frequency of SILP CD4+ T cells expressing Ki67, GzB, and IFN𝛾 from SPF and GF mice. (E-F) Representative flow cytometry plots showing (E) frequency of SILP CD4+ RORgt+ T cells and (F) the frequency expressing IL-17a from SPF and GF mice. (G) Uniform Manifold Approximation and Projection (UMAP) plot of SILP CD4 T cells split by condition. Clusters are annotated into 11 cell types. (H) Single cell expression of *Tbx21* (T-bet) projected on the UMAP plot of SILP CD4 T cells split by condition. Cells where *Tbx21* expression is not detected are shown in gray. (I) Single cell expression of the iNKT chain rearrangement TRAV11-TRAJ18 projected on the UMAP plot of SILP CD4 T cells. Cells with other TRA rearrangements shown in blue, cells where no chain was detected are shown in gray. (J) Single cell expression of *Maf* projected on the UMAP plot of SILP CD4 T cells split by condition. Cells where *Maf* expression is not detected are shown in gray. (K) Single cell expression of *Ahr* projected on the UMAP plot of SILP CD4 T cells split by condition. Cells where *Ahr* expression is not detected are shown in gray. (L) Dot plot showing the normalized expression of the indicated genes in the indicated cell types. Dot size represents the faction of the cells where the gene is detected, and the color scale represents average normalized expression. (B - F) Gated on CD45+ CD90.2+ TCRb+ CD4+ T cells. Data are representative of at least three independent experiments (A - F). Ns, not significant, *P < 0.0332, **P < 0.0021, ***P < 0.0002, ****P < 0.0001. Multiple unpaired t tests with Welch correction. Error bars indicate SEM.

**Figure S3:**
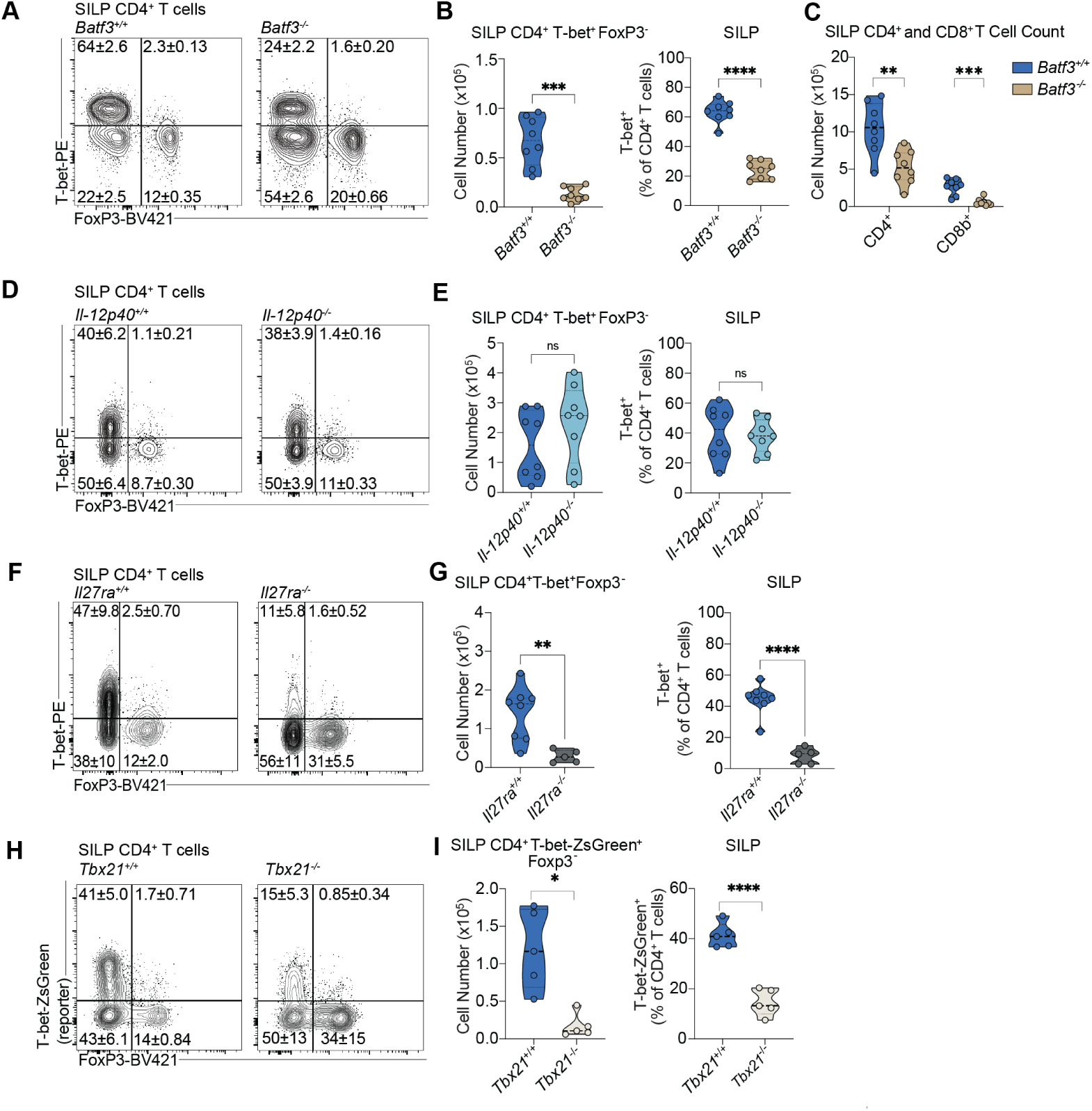
IL-27 and cDC1 factors, but not IL-12, control the accumulation of SILP CD4⁺T-bet⁺ T cells. (A) Representative flow plots showing T helper frequencies on gated CD4 T cells between co- housed WT and *Batf3^-/-^* mice at homeostasis. (B) Violin plots of SILP Tbet+ Foxp3- CD4 T cell counts and frequencies depicted in (A), respectively. (C) Violin plots off of SILP total CD4+ and CD8+ T Cell counts between cohoused WT and *Batf3^-/-^* mice at homeostasis. (D) Representative flow plots showing T helper frequencies on gated CD4 T cells between cohoused WT and *Il-12p40⁻* mice at homeostasis. (E) Violin plots of SILP Tbet+ Foxp3- CD4 T cell counts and frequencies depicted in (D), respectively. (F) Representative flow plots showing T helper frequencies on gated CD4 T cells between cohoused WT and *Il27ra^-/-^* mice at homeostasis. (G) Quantification of SILP Tbet+ Foxp3- CD4 T cell counts and frequencies depicted in (F), respectively. (H) Representative flow plots showing SILP CD4+ T-bet+ frequencies between T-bet-ZsGreen and T-bet-ZsGreen-*Tbx21^-/-^* mice at homeostasis. (I) Quantification of SILP Tbet+ Foxp3- CD4 T cell counts and frequencies depicted in (H), respectively. All flow plots are gated on CD45+ CD90.2+ TCRb+ Cd1d- CD4+ T cells. All violin plots, excluding (C), are gated on CD45+ CD90.2+ TCRb+ Cd1d- CD4+ T cells. Data are representative of at least three independent experiments. Ns, not significant, *P < 0.0332, **P < 0.0021, ***P < 0.0002, ****P < 0.0001. Comparisons using Mann-Whitney test (C, E, G, I) or Welch’s t test (B). Error bars indicate SEM.

**Figure S4:**
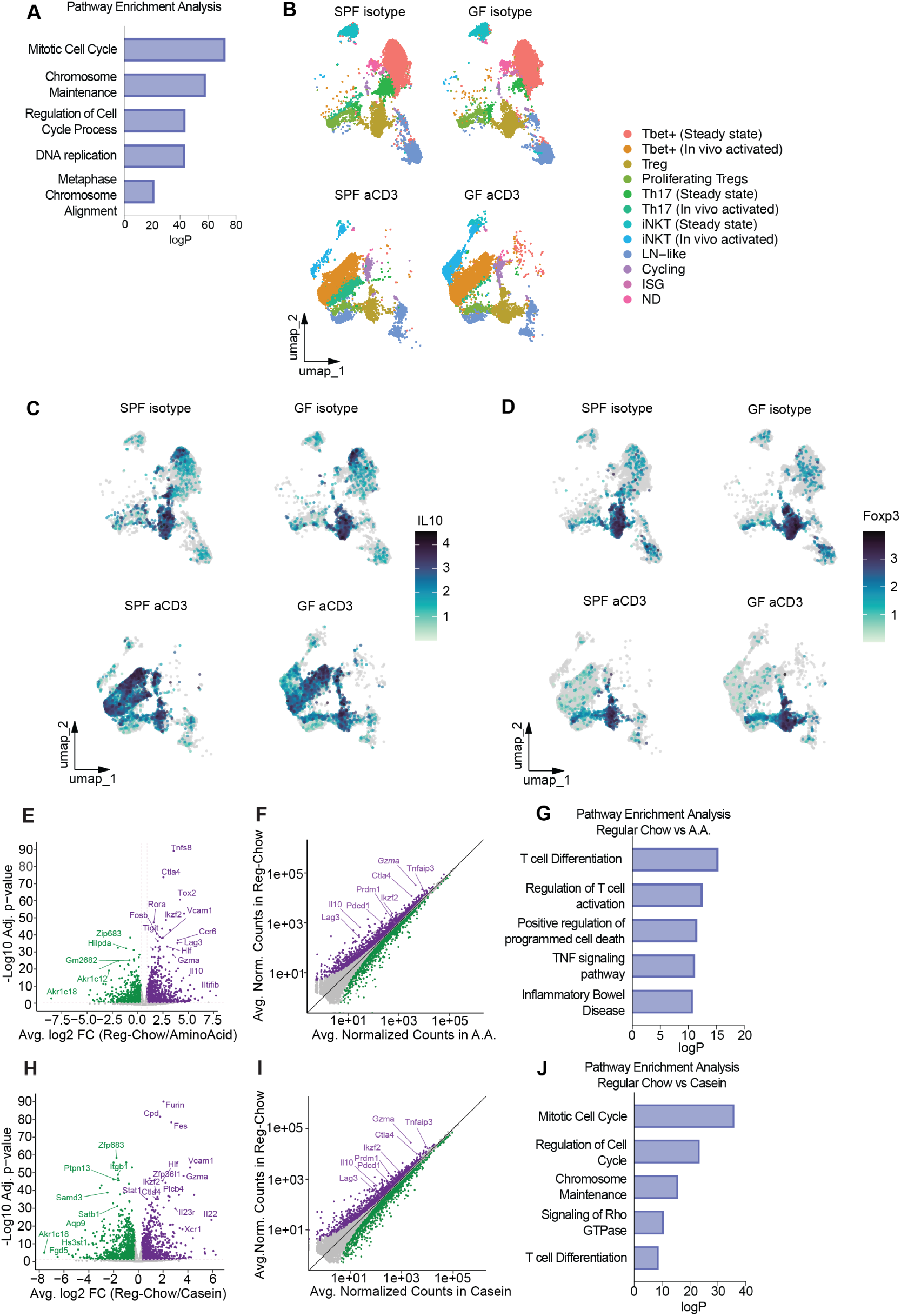
related to Figure 4: In vivo activation reveals a suppressive signature and IL-10 expression capacity of small intestinal CD4⁺T-bet⁺ T cells. (A) Pathway enrichment analysis showing significant GO terms from genes significantly upreg- ulated by ⍺CD3 injection of the bulk RNA sequencing experiment shown in Figure 4 A-D. (E-J) Single Cell RNA sequencing of sorted SILP CD4⁺ T cells from SPF and GF mice injected with isotype or ⍺CD3 antibody 24 hours prior to tissue collection (3-4 mice per group). (B) Uniform Manifold Approximation and Projection (UMAP) plot of SILP CD4⁺ T cells from ⍺CD3- vs isotype-injected mice split by condition. Clusters are annotated into 12 cell types. (B) Single cell expression of *Il10* projected on the UMAP plot of SILP CD4⁺ T cells from ⍺CD3- vs isotype-injected mice split by condition. Cells where *Il10* expression is not detected are shown in gray. (D) Single cell expression of *Foxp3* projected on the UMAP plot of SILP CD4⁺ T cells from ⍺CD3- vs isotype-injected mice split by condition. Cells where *Il10* expression is not detected are shown in gray. (E-G) Pseudobulk analysis of Single Cell RNA sequencing data of SILP CD4⁺Tbet⁺Foxp3⁻ T cells from SPF mice. (E) Volcano Plot showing differentially expressed genes in sorted SILP CD4⁺Tbet⁺Foxp3⁻ T cells from SPF ⍺CD3- vs isotype-injected mice as measured by pseudobulk analysis of single cell RNAseq data. (F) Scatter Plot showing differentially expressed genes in sorted SILP CD4⁺Tbet⁺Foxp3⁻ T cells from SPF ⍺CD3- vs isotype-injected mice as measured by pseudobulk analysis of single cell RNAseq data. Genes indicative of a suppressive signature are highlighted. (G) Pathway enrichment analysis showing significant GO terms from genes significantly upregulated by ⍺CD3 injection in SPF SILP CD4⁺Tbet⁺Foxp3⁻ T cells as measured by pseudobulk analysis of single cell RNAseq data. (H-J) Pseudobulk analysis of Single Cell RNA sequencing data of SILP CD4⁺Tbet⁺Foxp3⁻ T cells from GF mice. (H) Volcano Plot showing differentially expressed genes in sorted SILP CD4⁺Tbet⁺Foxp3⁻ T cells from GF ⍺CD3- vs isotype-injected mice as measured by pseudobulk analysis of single cell RNAseq data. (I) Scatter Plot showing differentially expressed genes in sorted SILP CD4⁺Tbet⁺Foxp3⁻ T cells from GF ⍺CD3- vs isotype-injected mice as measured by pseudobulk analysis of single cell RNAseq data. Genes indicative of a suppressive signature are highlighted. (J) Pathway enrichment analysis showing significant GO terms from genes significantly upregulated by ⍺CD3 injection in GF SILP CD4⁺Tbet⁺Foxp3⁻ T cells as measured by pseudobulk analysis of single cell RNAseq data. Error bars indicate SEM.

**Supplementary table 1:**
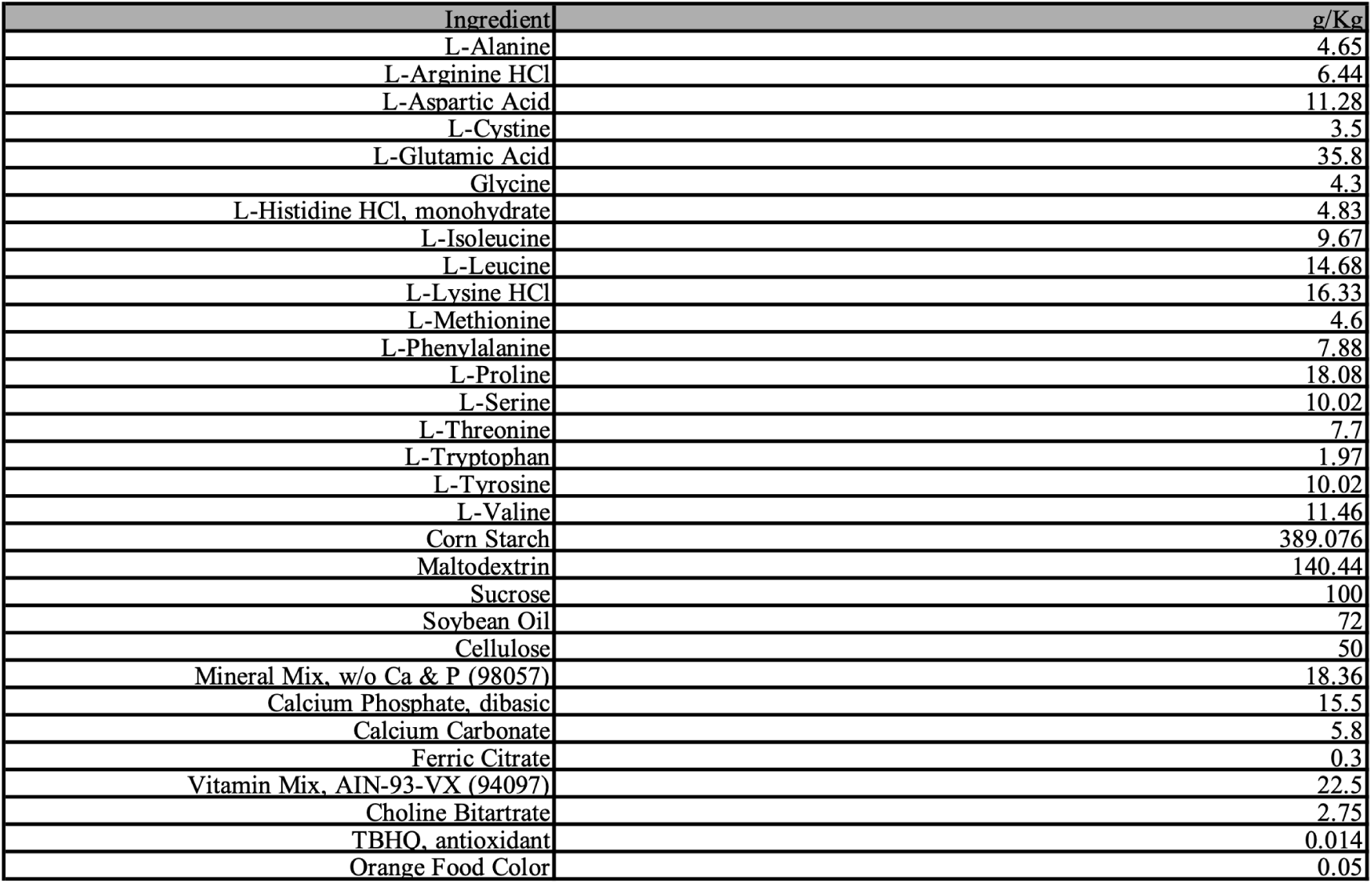
Amino Acid Diet Composition.

**Supplementary table 2:** Casein Diet Composition.

| Ingredient | g/Kg |
| --- | --- |
| Casein | 200 |
| L-Cystine | 3 |
| Corn Starch | 371.336 |
| Maltodextrin | 132 |
| Sucrose | 100 |
| Soybean Oil | 70 |
| Cellulose | 50 |
| Mineral Mix, AIN-93G-MX (94046) | 48 |
| Ferric Citrate | 0.3 |
| Vitamin Mix, AIN-93-VX (94047) | 22.5 |
| Choline Bitartrate | 2.75 |
| TBHQ, antioxidant | 0.014 |
| Red Food Color | 0.1 |

